# Neuromuscular impairments alter energetic cost landscape curvature and stride speed variability in post-stroke locomotion

**DOI:** 10.64898/2026.04.27.721024

**Authors:** Miles Smith, Praneeth Namburi, Nidhi Seethapathi, Brian W. Anthony

## Abstract

Neuromuscular impairments induce compensatory effects which alter the dynamics of human movement, but the mechanism linking specific impairments to post-stroke locomotion remains poorly understood. Here, we combine a predictive neuromusculoskeletal simulation framework with experimental gait observations in stroke survivors to test how two hemiparetic impairments, reduced muscle strength and increased baseline muscle activity, reshape the energetic cost landscape. We then evaluate whether impairment-dependent changes in the cost landscape curvature are associated with stride speed variability, which is experimentally observed to be higher after stroke. Using neuromusculoskeletal simulations, we show that increased paretic muscle activity reduces local curvature near the cost-minimized speed more than reduced paretic muscle strength and find that this predicts increases in stride speed variability observed in hemiparetic locomotion. These results support a mechanistic hypothesis that flatter cost landscapes reduce the relative cost of suboptimal behavior and, therefore, may contribute to increased motor variability after stroke.

**Author summary:** Motor disorders, such as those caused by stroke, change the way humans walk and interact with the world, often reducing quality of life. Stroke affects millions of people each year and commonly causes unilateral motor impairment that requires substantial rehabilitation. In this study, we use simulations to understand how different impairments, specifically reduced muscle strength and increased muscle activity, alter movement behavior. By varying the severity of the impairment and the speed of walking, we show how different impairments lead to compensation strategies and influence sensitivity to speed fluctuations. These findings suggest that the energetic penalty incurred from suboptimal behavior changes with neuromuscular impairment and are predicted to contribute to increased variability after stroke.

## Introduction

Human movement is often modeled as the outcome of an optimization process, in which motor behavior emerges as a trade-off between energetic cost and reliable performance of the task [1–5]. Neuromuscular impairments can alter this optimization by imposing additional constraints and inducing compensation strategies resulting in altered muscle mechanics and coordination patterns, and restructuring of the energetic cost landscape [6–9]. Establishing mechanistic links between neuromuscular impairments and movement performance through experiments alone is challenging because physiological parameters cannot be independently controlled *in vivo*. Predictive neuromusculoskeletal simulations provide insight by linking muscle-tendon mechanics to emergent gait patterns and the cost of transport during locomotion [10–12]. Bridging simulations and observations can deepen the mechanistic understanding of impairment effects and support the development of personalized rehabilitation strategies, physiologically-informed assistive devices, and computational biomarkers that can be used for health monitoring and diagnostics [13–16].

Due to the implementation of direct collocation methods, predictive neuromusculoskeletal simulations based on trajectory optimization have enabled the prediction of emergent gait behavior based on optimality principles [12, 17, 18]. This introduces methods that can be used to predict the energetic cost landscapes under a set of neuromusculoskeletal constraints and speeds given a physiologically-inspired objective function [13]. These simulations have been used to identify physiological mechanisms that underlie changes in the cost of transport and optimal walking speed in impaired and aged locomotion [11, 13, 14, 19, 20]. While neuromechanical models have been used to understand stability and variability in locomotion based on kinematic variability, neuromusculoskeletal models have rarely been used to gain insight into the structure of variability from an energetic perspective despite theoretical and behavioral work suggesting that variability can reflect sensitivity to changes in cost [21–26]. This work examines how neuromuscular impairments reshape the cost landscape and, specifically, how impairment-driven changes in the cost of deviating from the optimal walking speed are associated with stride speed variability.

The energetic cost landscape of locomotion can be characterized by three features: the speed at which the cost of transport is minimized, called the optimal walking speed, the energetic cost of transport at that optimal point, and the curvature of the landscape around the optimum, which describes how steeply the cost increases from deviations from the optimal speed [8, 25, 27, 28]. In this framework, compensatory effects, such as changes in optimal speed, energetic cost, and muscle-level effort distribution, emerge from impairment-imposed neuromuscular constraints. Here, we test whether impairment-dependent changes in the cost landscape curvature are associated with stride speed variability across gait cycles (Fig. 1). Thus far, most studies on predictive neuromusculoskeletal simulations have focused on changes in optimal speed or energetic cost of transport, but discussions on the energetic penalty for suboptimal behavior described by the curvature of the cost landscape are relatively unexplored [13, 14, 29–32]. Specifically, it is hypothesized that a flatter cost landscape means that deviations from the optimal locomotion speed incur relatively small energetic penalties, reducing the system’s resistance to changes in speed and increasing the expected variability in speed between strides.

**Fig 1.**
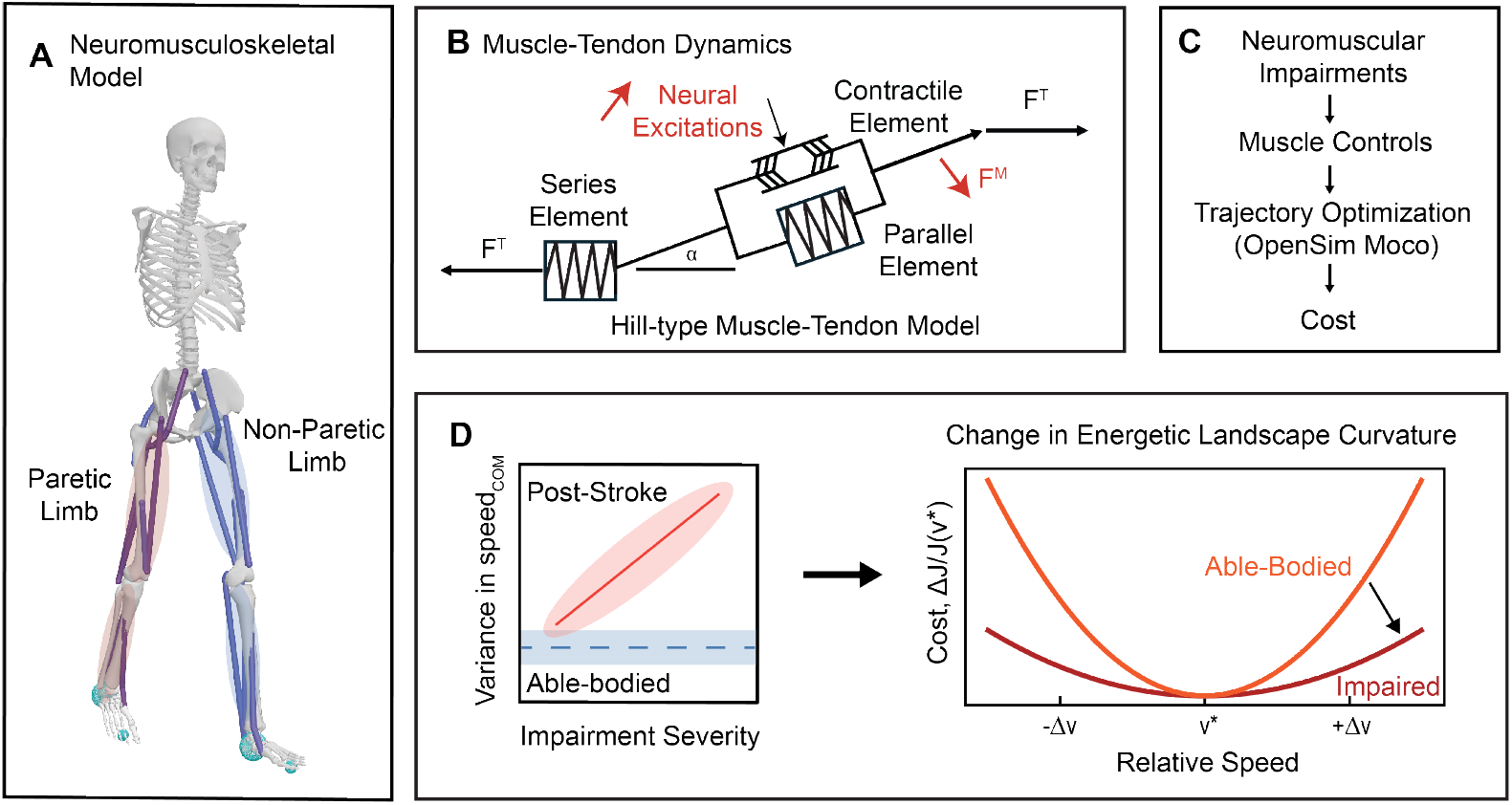
Optimization-based predictive simulations were used to impose neuromuscular impairments on a 2D gait model and assess how impairments influence changes in cost during pathological locomotion. (A) Hemiparetic locomotion was simulated using a 2D gait model. Neuromuscular impairments were imposed on the paretic limb, while the non-paretic limb was not altered. (B) Muscle-tendon dynamics are captured by a Hill-type muscle-tendon model. Constraints were imposed on the neural excitations to the paretic limb and the strength of the muscles in the paretic limb. (C) The cost of transport is resolved through trajectory optimization given the neuromuscular impairments and speed goal. (D) It is hypothesized that changes in the curvature of the cost landscape, a descriptor of cost sensitivity, are related to experimental observations of motor variability. Specifically, it is proposed that reducing the cost sensitivity will increase the observed variability.

To assess this hypothesis, we focus on hemiparetic locomotion observed after stroke and examine two key neuromuscular impairments observed in chronic hemiparesis: increased muscle activity and reduced muscle strength in the paretic limb, and examine how these impairments influence the cost landscape [7, 33–37]. Post-stroke hemiparetic gait is characterized by slower preferred walking speeds, increased metabolic cost, asymmetries between the paretic and non-paretic limb, and increased spatiotemporal stride variability [38–42]. Although increased spatiotemporal gait variability has been associated with an increased risk of falls in older adults and clinical populations, it is not clear how specific neuromuscular impairments influence gait variability due to the inherent difficulty in experimentally quantifying the effect of isolated impairments on coordinated movement [43–46].

Using a predictive neuromusculoskeletal simulation framework, we varied the speed of movement and severity of impairment in a two-dimensional musculoskeletal model to compute the cost of transport surfaces, which define the cost landscape during locomotion. We show that neuromuscular impairments reshape the cost landscape not only by changing the optimal movement strategy and energetic cost of transport, but also by altering the curvature of the cost landscape, the energetic penalty for deviating from optimal behavior. Impairment-driven reductions in the cost landscape curvature provide a mechanistic explanation for increased stride speed variability in post-stroke locomotion. These predictions are validated against gait observations in a post-stroke clinical population, where impairment severity and stride speed variability are significantly correlated, which is consistent with a relationship between neuromuscular impairment and motor variability mediated by the curvature of the energetic cost landscape. Reduced landscape curvature is observed more strongly in simulations with increased paretic muscle activity than in simulations with reduced paretic muscle strength, suggesting that the type of impairment can influence how the cost landscape is reshaped, not just the severity.

## Materials and methods

### Predictive neuromusculoskeletal simulations

To understand the effects of hemiparetic impairments, a framework was developed for predictive neuromusculoskeletal simulations using OpenSim Moco, which utilizes direct collocation-based trajectory optimization [10–12, 47]. The framework accepts two independent variables as inputs and minimizes a cost function through trajectory optimization to recover cost-optimal muscle activations throughout one stride (Fig. 1). For each simulation, the initial conditions and model were kept constant, while the gait speed and the severity of the impairment were manipulated. A key aim of this framework is to establish a methodology that captures subtle changes in kinematics, dynamics, and energetics from simulations. Then, by comparing the relative changes, more generalizable links between impairment, speed, and locomotion can be identified that smooth over the inaccuracies of any individual model or simulation. An added benefit of automating the optimization process is that the total computational time decreases because optimizations can be performed distributed among the available resources [48].

Two different sets of simulations were developed to capture the isolated effects of the impairments commonly observed in stroke survivors, (1) reductions in muscle strength and (2) increases in muscle activity [33, 34, 37, 49–52]. A 2D gait model with 10 muscles, 18 degrees-of-freedom, and 4 contact objects was chosen for these simulations because of its computational simplicity, which allows for relatively quick iteration and optimization (Fig. 1A). Hemiparetic impairments were imposed directly on the muscles on the right side of the model, known as the paretic limb, with uniform severity in the muscles on the paretic side. No impairments were imposed on the left side of the model, referred to as the non-paretic limb, to allow for unilateral compensation strategies to emerge.

Neuromuscular impairments were imposed by reducing the *MaxIsometricForce* of the muscles on the paretic side of the model or increasing *MinControl* of the muscles on the paretic side of the model (Fig. 1B). Elevated paretic baseline muscle activity was modeled by increasing *MinControl* because it sets a lower bound on neural excitation, which prevents full muscle relaxation during the gait cycle. This would most closely represent tonic baseline excitations that are observed clinically post-stroke [33, 49–51, 53, 54]. Reduced paretic muscle strength was modeled by scaling down *MaxIsometricForce* in the paretic muscles, which reduces the maximum force each muscle can produce to model the effects of post-stroke muscle weakness [52, 55]. For each set of simulations, the unique pairings of impairment severity and goal gait speed for the simulation were uniformly distributed throughout the simulation space. In simulations with increased *MinControl*, 297 out of 340 simulations converged on solutions that met the constraint tolerance of 10^−4^ within 2000 iterations. In simulations with reduced *MaxIsometricForce*, 300 out of 339 simulations converged on solutions with the same criterion. The simulations were performed across batches that were parallelized across three tasks that each have 32 CPU cores with 4-GB of RAM. Optimizations that did not converge were flagged and removed from the analysis.

In each simulation, the optimizer aimed to primarily minimize the cumulative sum of muscle activations cubed during a 0.92 second simulation while also minimizing the difference between the goal velocity and the simulation velocity and the difference in joint positions between the initial and final time steps to capture a full stride (Eq. 1).

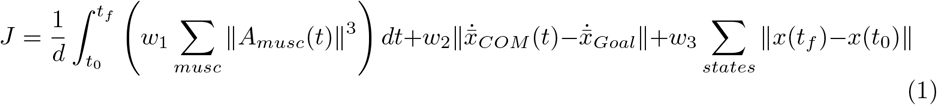

Cubed muscle activations are used in optimization as a proxy for metabolic cost because it has been shown to predict physiologically realistic muscle force sharing [3, 56, 57]. The time interval corresponds to approximately one gait cycle at moderate speeds, but at more extreme speeds this would influence the stride length. All cost-function weights were held constant and equal throughout the simulation space.

### Cost of transport

For comparison, the cost of transport was used to compare the simulations in the simulation space. Two definitions for the cost of transport were considered. First, the fatigue cost of transport (FOT), which corresponds to the muscle activation term in the cost function, or the minimized form of the cost function [14, 18, 58]

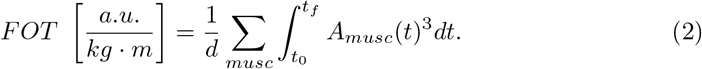

Second, the metabolic cost of transport (COT) was estimated using Bhargava’s model for metabolic energy consumption based on heat generation and mechanical work for each muscle,

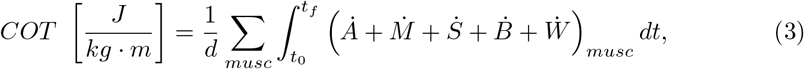

where 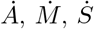, and 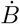 represent the heat generation rates of muscle activations, activation maintenance, muscle shortening, and the basal metabolic rate, respectively, and 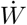 is the mechanical work rate of muscle contraction [59]. We compute both system-level and muscle-level costs, to construct the cost landscapes across both scales. Each energetic model was applied across the complete system and for individual muscles to characterize the interplay between energy requirements across the system and between individual muscles.

Because FOT defines the objective function that governs predictive simulations, it represents the effective cost landscape that shapes optimized behaviors. It is important to note that this creates a degree of circularity because FOT is the quantity the optimizer minimizes. The resulting cost landscape reflects the model’s optimization target rather than a directly measured physiological quantity. Conclusions drawn from the FOT curvature are therefore model internal, rather than direct evidence of the quantity that humans optimize during locomotion. Evaluating both FOT and COT allows for a distinction between the optimization-based FOT landscape and the estimated physiological metabolic cost landscape. Each energetic model was applied at the system and individual muscle levels to characterize how energetic demands are distributed across scales, but FOT landscapes produce cost-minimized speeds that are more closely aligned with the preferred walking speeds of stroke survivors. This discrepancy between the cost-minimized speed in the FOT and COT landscapes also limits the relevance of COT curvature estimates for characterizing post-stroke locomotion.

### Curvature of the cost landscape

Stride variability in walking speed was quantified from experimental gait data, but was not simulated within the predictive modeling framework. Instead, simulated energetic landscapes were used to estimate the curvature, which was then tested as a mechanical predictor of the empirically observed variability. Motivated by the proportional sensitivity observed in biological systems, it is assumed that the motor control system is sensitive to relative changes in cost [28, 60]. This motivates a comparison of the cost landscape relative to the minimum cost for a given state of impairment. To estimate the curvature of the cost landscape, the cost landscape is approximated to be a second-order Taylor series expansion about the optimal point to estimate the local curvature of the landscape. This is formalized as,

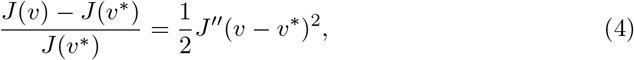

where *v* is the speed of locomotion, *v*^∗^ is the cost-minimized locomotion speed, *J* (*v*) is the cost at a given speed, and *J* ^′′^ is the curvature of the cost landscape relative to the cost-minimized speed. The curvature of the cost landscape is descriptive of the cost sensitivity to deviations from the optimal speed. The curvature was derived from the fatigue cost of transport landscape (Eq. 2), which captures muscle activations in the model during a complete stride in steady-state locomotion.

Simulations with *MinControl* greater than 0.7 were excluded from the local curvature analysis because of the introduction of a non-physical walk-to-run transition. Similarly, simulations with *MaxIsometricForce* less than 0.4 were excluded as this exceeds the range of paretic weakness observed in ambulatory stroke survivors, where inter-limb muscle strength asymmetries have been shown to be up to 60% [61].

### Relationship between cost landscape curvature and stride speed variability

Assuming that fluctuations in stride speed follow a Boltzmann-like distribution and relative changes in cost can be expanded as a second-order Taylor series, it is expected that stride speed variability would follow a Gaussian distribution where the coefficient of variation (CV) is described as (S1 Appendix),

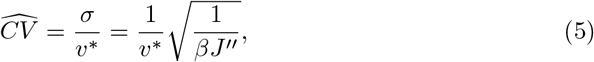

where *σ* is the standard deviation of the stride speeds, *v*^∗^ is the optimal walking speed, *J* ^′′^ is the local relative curvature of the cost landscape, and *β* is a scaling factor. To estimate the coefficient of variation from the curvature of the cost landscape, *β* was estimated by comparing the cost landscape curvature without impairment with the average coefficient of variation in the able-bodied population, such that

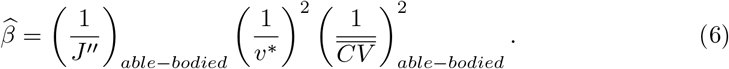

### Non-paretic propulsion

The fraction of non-paretic propulsion (*P*_*NP*_) was estimated in simulation as a means of comparing the impulses generated during a stride in the non-paretic limb to the paretic limb. The impulse of a limb was estimated based on the ground reaction forces during the simulated gait cycle, such that

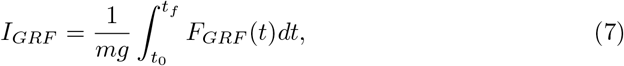

where *I*_*GRF*_ is the impulse generated through ground contact, *m* is the subject mass, *g* is acceleration due to gravity, *t* is time, and *F*_*GRF*_ is the ground reaction forces from foot contact. The fraction of non-paretic propulsion is defined as,

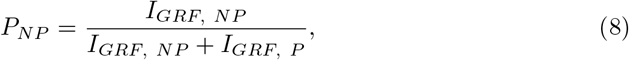

which compares the impulses generated by the non-paretic limb (NP) to the impulses generated between both the paretic (P) and non-paretic limbs. In symmetric locomotion non-paretic propulsion should be 0.5. Values greater than 0.5 would indicate proportionally higher impulses from the non-paretic limb than the paretic limb.

### Experimental observations

We analyzed a publicly available dataset comprising 138 able-bodied adults and 50 stroke survivors to estimate stride speed variability and physiological activity as the subjects walked along a 12-m walkway [62]. From this dataset, kinematic data from marker-based motion capture data was collected for each subject as they walked across the walkway. Surface electromyography (EMG) data was also collected from key muscles in both the paretic and non-paretic limbs, such as the gastrocnemius, a calf muscle, and the rectus femoris, a quadricep muscle. The EMG data was processed to capture the EMG activity of the individual muscles in a gait cycle. Full details of the EMG filtering, rectification, and normalization pipeline are described in Van Criekinge et al. [62]. For comparison between subjects, the muscle activity is resolved to the cumulative muscle activity normalized by the duration of the gait cycle.

K-means clustering was used to split the group of stroke survivors into a group with mild and high muscle activity in the paretic limb based on the muscle activity in the paretic rectus femoris measured during a gait cycle. The rectus femoris was selected as the variable for clustering because large changes in EMG activity have been observed in this muscle during post-stroke locomotion [33]. The gastrocnemius, which also shows post-stroke changes, was reserved as the primary outcome variable for validating the predicted compensation strategies. The population was split into two groups (*k* = 2) to produce groups that are directly comparable to the simulated impairment conditions, which compare mild and severe impairments in muscle activity.

Based on simulated observations and prior experimental observations, we associate the preferred walking speed of subjects with the severity of the impairment (S1 Fig) [35, 63]. To compare observations to the preferred walking speed of subjects, we estimate the Froude velocity (Fr), a characteristic velocity scale that normalizes speed by leg length, such that

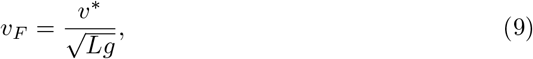

where *v*^∗^ is the preferred walking speed of the subject, *L* is their leg length, and *g* is acceleration due to gravity [64]. The typical Froude velocity for an able-bodied subject is approximately 0.42 [65].

### Variability in stride speed

Variations in stride speed were quantified using the coefficient of variation (CV), which scales the magnitude of the standard deviation by the mean velocity such that

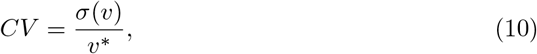

where *σ*(*v*) is the standard deviation of the stride speed. The coefficient of variation was chosen to compare stride speed variability between subjects since it normalizes the total variability by the expected speed to assess the relative magnitude of variability. The variance in stride speed was estimated using kinematic observations based on the time between heel contact points and the displacement in the anterior-posterior plane measured from the seventh cervical vertebrae between the heel contact points. The coordinate system of the motion capture system was rotated to align with the principal component of movement to estimate the displacement in the anterior-posterior plane. The stride speed variance was calculated for each subject with at least six computed stride speeds to balance inclusion across subjects with statistical power, consistent with prior spatiotemporal variability studies [40].

Differences in the coefficient of variation attributed to changes in speed were characterized using the scaling relationship observed in the able-bodied population and extrapolated across the population of both able-bodied individuals and stroke survivors, such that

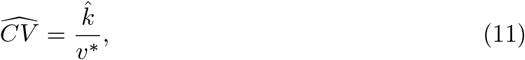

where 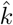 is an empirically derived parameter from the able-bodied population that was determined through least-squares regression. The residuals relative to the estimated coefficient of variation were estimated for each subject as

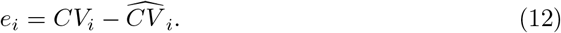

## Results

To test whether impairment-driven changes in cost landscape curvature explain increases in stride speed variability, experimental observations of stride speed variability are compared with cost landscapes produced from predictive neuromusculoskeletal simulations. Simulations reproduce known impairment-driven reductions in optimal walking speed and increases in cost of transport, and are extended to demonstrate that increased paretic muscle activity, but not reduced paretic muscle strength, reduce the local curvature of the cost landscape. The differences in curvature are explained by asymmetric non-paretic compensation strategies that emerge in simulations with increased paretic muscle activity. Applying a theoretical relationship between curvature and variability, the predicted coefficient of variation increases with the severity of muscle activity impairment, consistent with the relationship between clinical function mobility scores and observed stride speed variability in stroke survivors.

### Impaired motor control increases stride speed variability in post-stroke locomotion

Stroke survivors exhibit stride speed variability that exceeds what would be predicted from reductions in walking speed alone. The variability of stride, quantified as the coefficient of variation (CV) in stride speed, is inversely correlated with the preferred walking speed in both the able-bodied and stroke survivor populations (Fig. 2A). The scaling relationship between speed and CV in the able-bodied population (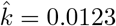, *R*^2^ = 0.27) was extrapolated to all subjects to assess the baseline contribution of speed to normalized variability. The modest fit in the able-bodied population is expected, as self-selected walking speeds in healthy adults produce a relatively uniform CV with limited speed-driven variability. The residual CV remained significantly elevated in stroke survivors compared to the able-bodied population (stroke survivor median residual = 0.021, able-bodied median residual = -0.006, Cohen’s *d* = 0.64, Welch t-test: t = 2.57, p = 0.015). This indicates that reduced walking speed alone does not explain increased variability after stroke despite reduced walking speed being a proxy of impairment associated with underlying neuromuscular changes (S1 Fig) [35]. The observation of an increased coefficient of variation in stride speed is consistent with previous observations reporting increased variability in stride time and step length after stroke [39, 40, 42].

**Fig 2.**
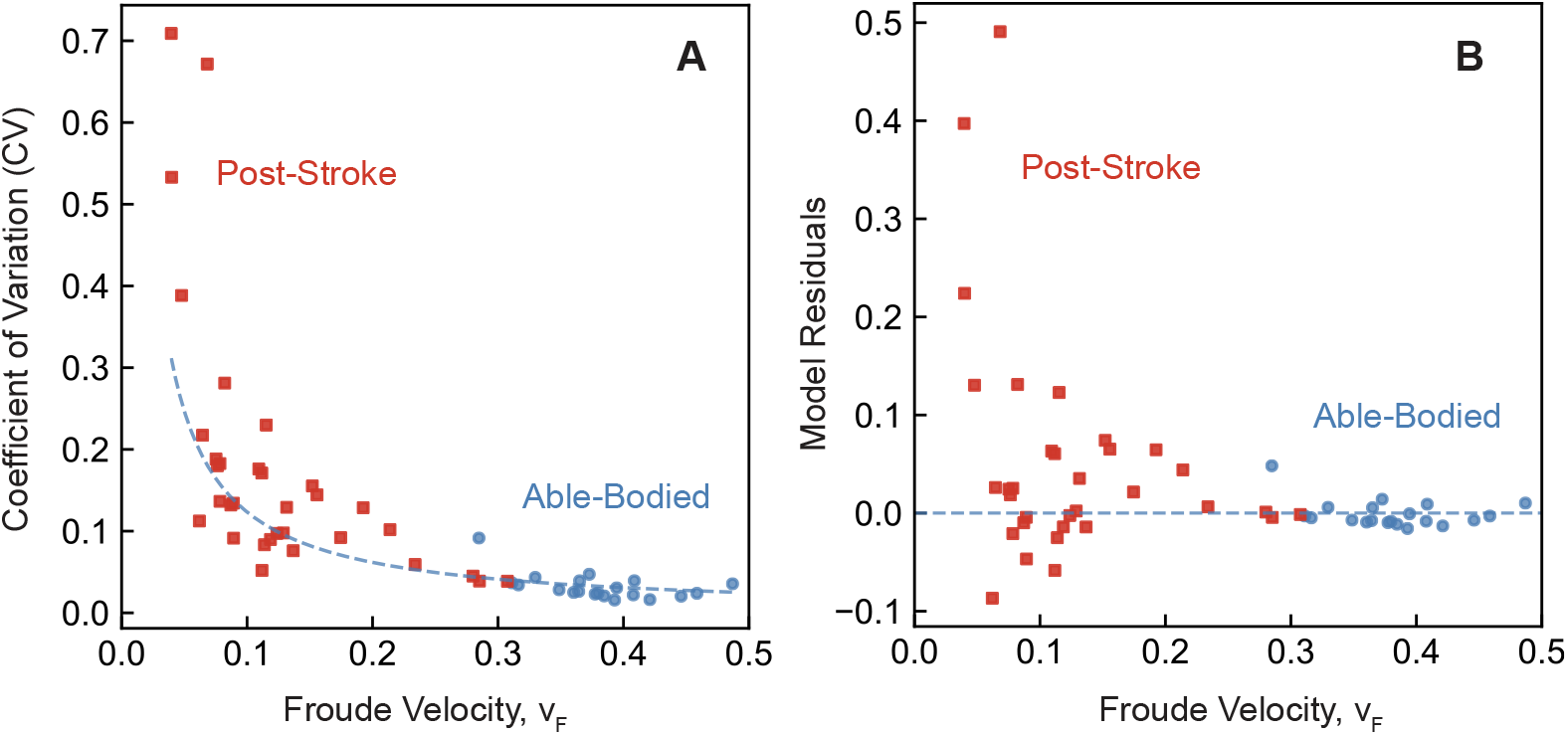
Stroke survivors show increased motor variability after controlling for changes in speed. (A) Both the post-stroke and able-bodied group are observed to be correlated with speed with a high statistical significance. Extrapolating the scaling between speed and CV in the able-bodied population (blue) to the population of stroke survivors, the coefficient of variation in the post-stroke population remains elevated relative to the extrapolated observations. (B) Residual variability among stroke survivors relative to the scaling relationship is significantly greater than the able-bodied population (Welch t-test, p=0.015), suggesting that reduced walking speed alone does not explain increased stride speed variability after impairment.

### Optimal behavior in neuromusculoskeletal simulations predicts impairment-driven reductions in walking speed and increases in cost of transport

Both reduced muscle strength and increased paretic muscle activity reduce the cost-minimized walking speed and increase the overall cost of transport in simulation. This is observed when considering optimal behavior in both the objective cost landscape (S2 Fig) and the metabolic cost landscape (S3 Fig). This aligns with previously reported clinical observations that neuromuscular impairments result in slower and more laborious locomotion [13, 35, 66]. In optimal behavior, it is observed that increased paretic muscle activity has a larger effect on both the fatigue and metabolic costs of transport as impairment severity increases (S2 Fig(C)) [14, 59].

Changes in both muscle strength and muscle activity influence optimized locomotion speed where the cost of transport is minimized. The fatigue cost of transport predicts optimal speeds at Froude velocity values between approximately 0.1 and 0.4 (S2 Fig(C)).

This is consistent with previous observations in predictive neuromechanical simulations [13, 14, 67]. When considering the metabolic cost of transport, the cost-minimized Froude velocities ranged between 0.4 and 0.6 (S3 Fig) [59]. Compared to the population of stroke survivors, which have a preferred Froude velocity between 0.05 and 0.3, more realistic cost-minimized locomotion speeds are observed when using fatigue cost of transport rather than a metabolic model to determine cost-minimized speed, motivating the use of the objective-aligned fatigue cost of transport over the metabolic cost of transport for analysis (Fig. 2) [14, 62].

### Increases in muscle activity reduce the local relative curvature of the cost landscape

Increased paretic muscle activity reduces the local curvature of the cost landscape, implying a reduced energetic penalty for deviating from the optimal speed. In contrast, reducing muscle strength in the paretic limb results in minimal changes in local curvature. Simulations with increased muscle activity in the paretic limb the curvature becomes smaller as the impairment severity increases, suggesting relatively small penalties in additional cost for suboptimal behavior compared to the overall level of fatigue (Fig. 3). In contrast, reductions in muscle strength in the paretic limb are not observed to significantly alter the local curvature despite reductions in muscle strength reducing the optimal walking speed in the simulations. This provides a cost-based explanation for the statistically observed increases in variability when assuming that subjects between groups are sensitive to the same relative changes in cost. The observations suggest that variations in stride speed are more strongly associated with changes in muscle activity in the paretic limb than changes in muscle strength in the paretic limb. To test this quantitatively, a theoretical relationship between curvature and variability was applied to predict the coefficient of variation from the simulated cost landscape (S1 Appendix).

**Fig 3.**
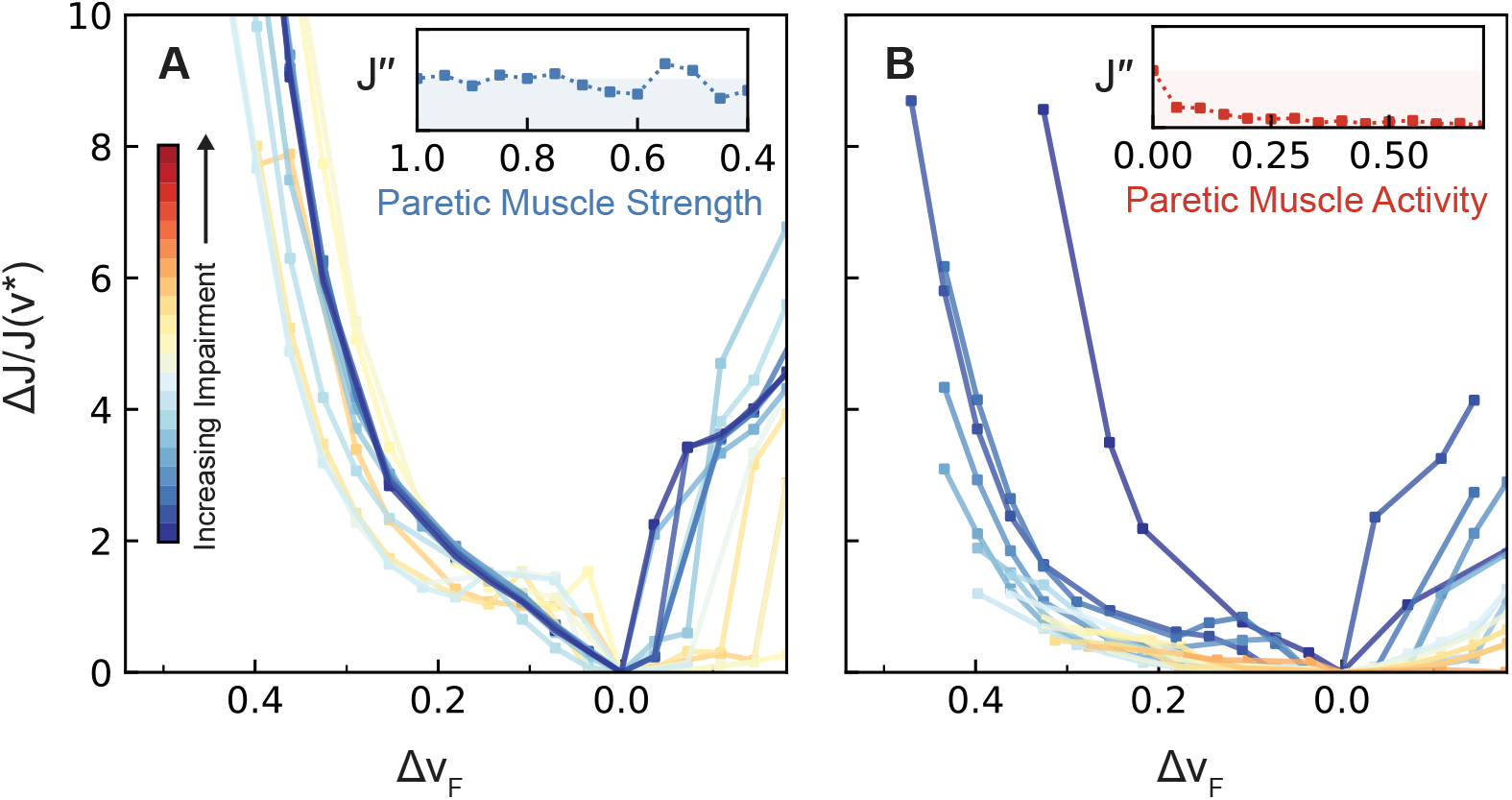
Increases in paretic muscle activity reduce the curvature of the cost landscape. (A) Reductions in muscle strength in the paretic limb result in movement behaviors that maintain a similar local curvature (*J* ^′′^) when compared against unimpaired behavior. (B) Increases in muscle activity in the paretic limb result in movement behaviors that reduce the local energetic curvature (*J* ^′′^) as the impairment severity increases.

### Asymmetric non-paretic compensation strategies drive reduced cost sensitivity under impaired muscle activity

The divergence in curvature between impairment types is explained by asymmetric non-paretic compensation strategies where increased muscle activity drives the cost-minimized solution into a regime of increased non-paretic propulsion, while reduced paretic muscle strength does not produce this shift. Throughout the cost landscape, compensation strategies emerge that create optimal movement strategies at different locomotion speeds and severity of impairment. These compensation strategies differ largely between the two impairment types, consistent with the observed changes in curvature (Fig. 3). Specifically, increased muscle activity in the paretic limb results in compensation strategies that are less sensitive to changes in cost as evidenced by the reduced landscape curvature. In contrast, reductions in muscle strength in the paretic limb result in compensation strategies that maintain a similar cost sensitivity, since the landscape curvature remains largely unchanged as impairment severity increases.

To understand why these two impairment types produce such different curvature responses, the difference in effort delivered between the paretic and non-paretic limb was assessed by estimating the fraction of non-paretic propulsion throughout the gait cycle (Eq. 8). When muscle strength is reduced, regions of high non-paretic compensation are observed in the cost landscape, but the cost-minimized speed avoids these asymmetric regions, maintaining similar propulsion strategies between limbs (Fig. 4D). In contrast, increasing paretic muscle activity forces the cost-minimized speed to introduce stronger asymmetric behaviors, with the non-paretic limb taking on a greater fraction of propulsion as impairment severity increases (Fig. 4A). This asymmetric compensation strategy may contribute to the mechanism through which increased paretic muscle activity could reduce cost landscape curvature, since the non-paretic limb’s increased contribution flattens the energetic penalty for deviations from the optimal speed.

**Fig 4.**
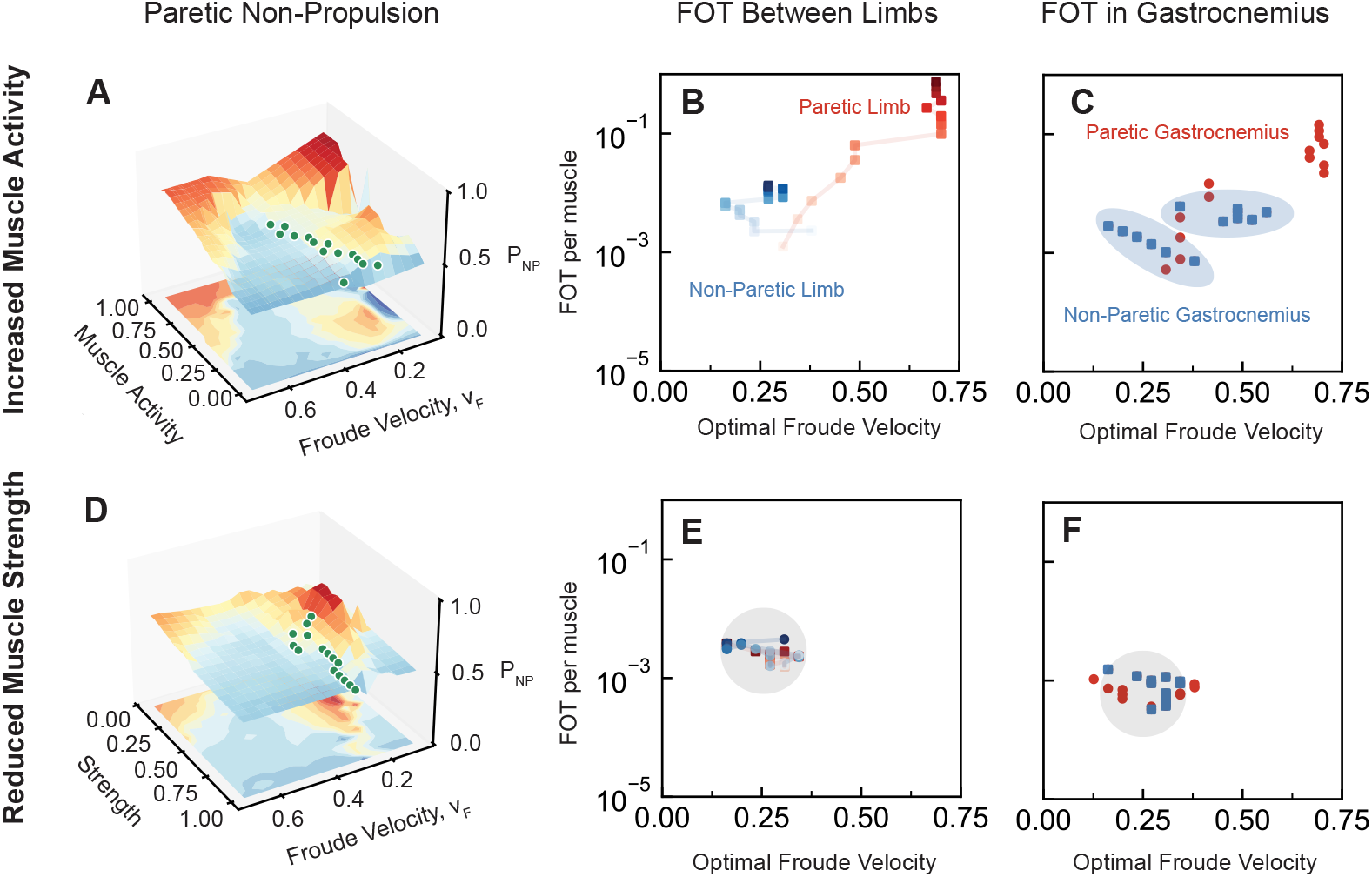
Increased paretic muscle activity is observed to shift the fraction of non-paretic propulsion to favor increased non-paretic drive while symmetric drive remains the optimal strategy as muscle strength is reduced. (A) Increased muscle activity in the paretic limb reveals a shift towards increased non-paretic propulsion from symmetric propulsion that the cost-minimized behavior (green) navigates through. This results in fatigue costs increasing with severity in the (B) non-paretic limb as a compensation strategy that is observed in the (C) gastrocnemius. (D) Reduced muscle strength results in a landscape where asymmetries become the favorable strategy, but the cost-minimized behavior (green) maintains similar propulsion strategies between the non-paretic and paretic limb. This results in similar fatigue costs between the (E) non-paretic and paretic limbs and (F) the non-paretic and paretic gastrocnemius.

Muscle-level compensation strategies in experimental observations and simulations were investigated to understand why changes in muscle activity alter curvature more than muscle weakness. In simulations with increased paretic muscle activity, non-paretic gastrocnemius is observed to increase despite slower cost-minimized speeds at high impairment severity, suggesting a muscle-level compensation effect observed due to increased paretic muscle activity (Fig. 4C). Two regimes are observed in the compensatory behavior of the non-paretic gastrocnemius, a milder and severe regime, in the non-paretic gastrocnemius when comparing the fatigue cost of the gastrocnemius to the muscle’s optimal locomotion speed. In the mild regime, muscle activity in the non-paretic gastrocnemius is negatively correlated with the optimal behavioral speed of the gastrocnemius, which suggests that increases in effort emerge to compensate for the impaired paretic limb. In the severe regime, gastrocnemius activity is positively correlated with optimal muscle behavior, suggesting a change in behavior, resulting in an increase in the cost-minimized speed for the individual muscle. In contrast, when reducing paretic muscle strength, symmetric strategies between the limbs and individual muscles remain preferable (Fig. 4 (E, F)).

The simulated prediction of increased non-paretic muscle activity in stroke survivors with increased paretic muscle activity was tested by comparing stroke survivors with the highest muscle activity in the paretic limb during locomotion and quantifying the muscle activity in the non-paretic gastrocnemius. Using k-means clustering to classify stroke survivors based on paretic muscle activity in the rectus femoris, a high muscle activity group was identified in the stroke survivor population (S4 Fig). A statistically significant negative correlation is observed between the preferred walking speed and muscle activity in the non-paretic gastrocnemius in stroke survivors of the high muscle activity group (S5 Fig). In contrast, no statistically significant relationship is observed between muscle activity in the non-paretic gastrocnemius and preferred walking speed in the able-bodied group or the remaining group of stroke survivors. The observed phenomenon of increased muscle activity in the compensating limb is consistent with previously reported findings of non-paretic compensation and is associated with asymmetries in mechanical work delivered by the paretic limb and non-paretic limb [68–70].

### Benchmarking of simulated muscle activation profiles against experimental EMG

Simulated compensatory gastrocnemius activations show moderate-to-good agreement with experimental EMG observations in stroke survivors (cross-correlation 0.7–0.8), while rectus femoris agreement is lower (0.4–0.5), consistent with the limitations of a 2D gait model (S6 Fig (C, F)). The simulated gastrocnemius and rectus femoris and the corresponding mean EMG activity during a gait cycle in the non-paretic limb at the optimal speed were compared in both the cases of impairment to assess the similarity of compensatory strategies between the simulations and the observed population (S6 Fig (A, B, D, E)). This was similarly observed in the paretic limb, where the gastrocnemius was found to have much higher cross correlation coefficients than the rectus femoris in both sets of impairments (S4 Fig). Slightly higher cross correlation coefficients were also observed in the paretic gastrocnemius (0.8-0.9) than in the non-paretic gastrocnemius (0.7-0.8). The low similarity of the rectus femoris is reasonable in a 2D gait simulation since the 3D effects observed in hemiparetic locomotion, such as hip circumduction, are not captured without muscles that can generate forces in the medio-lateral direction [70–72]. In contrast, the gastrocnemius is a key muscle in the anterior-posterior plane because it generates a significant amount of propulsive force and consumes a large fraction of the overall metabolic cost during locomotion, suggesting that behavior of the gastrocnemius should be captured with higher accuracy in a 2D gait model [69, 73].

### Cost landscape curvature provides a mechanism for impairment-driven increases in stride speed variability

Predicted stride speed variability increases with paretic muscle activity severity, consistent with the empirical correlation between stride speed variability and functional mobility scores in stroke survivors. To quantitatively link cost landscape curvature to stride speed variability, a theoretical relationship was derived assuming a Boltzmann-like distribution of stride speeds (Eq. 5, see S1 Appendix for full derivation). The relationship was applied by fitting a single scaling parameter, *β*, which scales the sensitivity of the sensorimotor system to changes in energetic cost. The scaling parameter was calibrated using the curvature of the cost landscape without neuromuscular impairments and the mean coefficient of variation from the able-bodied population and then applied across all impaired simulations to predict the coefficient of variation based on the impairment-driven changes in the cost landscape.

In the clinical dataset, stride speed coefficient of variation is significantly correlated with reported Performance Oriented Mobility Assessment (POMA) scores, providing evidence that high degrees of functional mobility impairment are associated with greater stride speed variability in sitroke survivors (Fig. 5(A)). Applying the theorized relationship between local curvature and CV, it is predicted that CV increases significantly in simulations with increased paretic muscle activity, but does not in simulations with reduced muscle strength (Fig. 5(B)) and aligns with the predictions for the cost landscape curvature (Fig. 3). The predicted CV when paretic muscle activity increases ranges from 3% to 14%. This underestimates the CV observed in the clinical population, which is linearly modeled to range between 3% and 33% depending on the impairment severity, but is calculated to be over 40% in some individuals. In contrast, reduced muscle strength predicts CV values between 2% and 4%, which is more in line with expected values in able-bodied subjects. The underestimation of CV is consistent with expectations for a model excluding sensorimotor noise and feedback, which would be expected to influence stride-to-stride fluctuations and contribute to variability beyond what the cost landscape curvature alone would predict.

**Fig 5.**
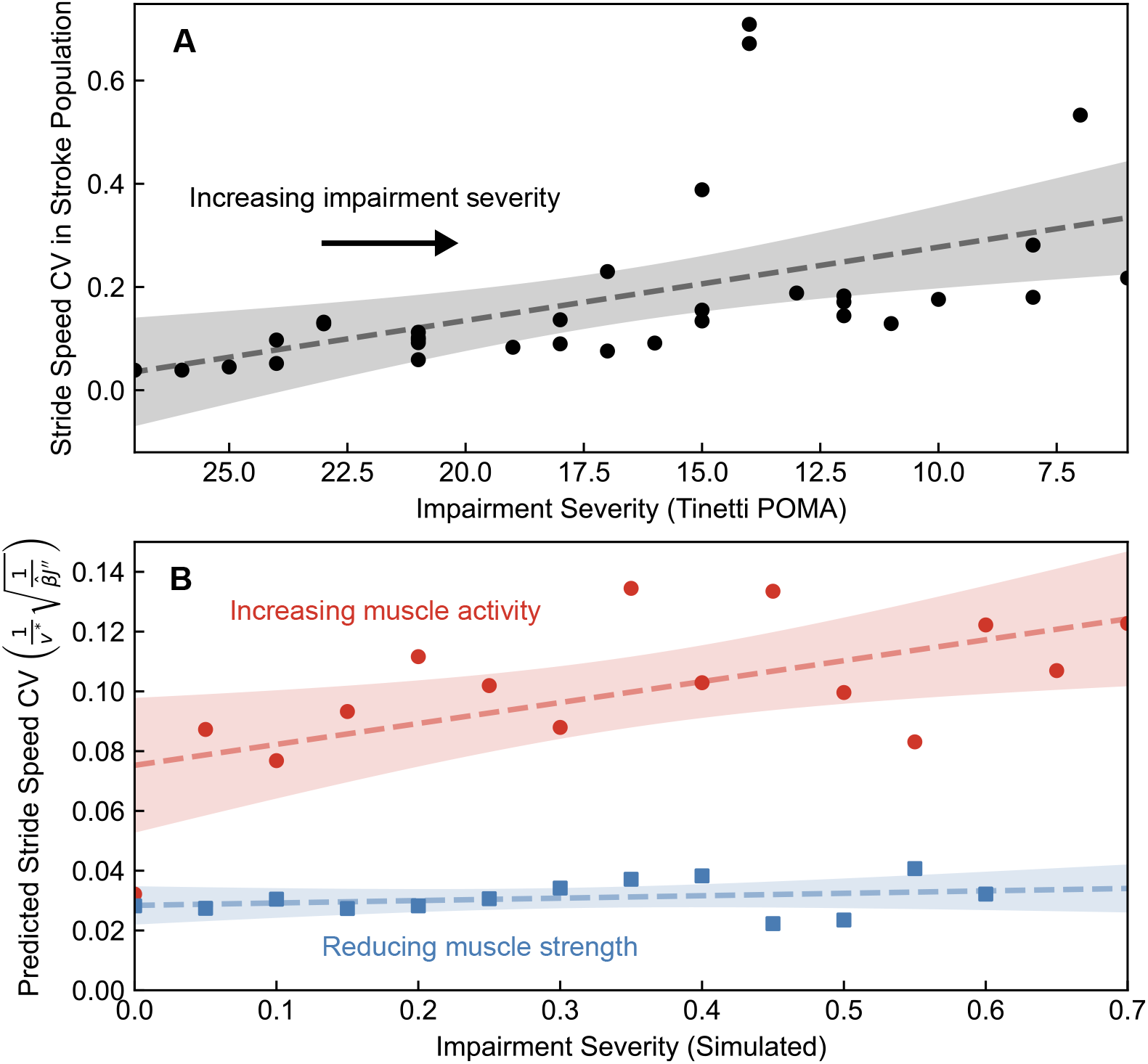
Stride speed coefficient of variation increases in clinical populations and simulations with increased muscle activity as impairment severity increases. (A) Stride speed CV is significantly correlated with Tinetti Performance Oriented Mobility Assessment (POMA) scores, a clinical measure of functional mobility and a proxy for impairment severity (Spearman *ρ* = −0.78, *p <* 0.001, *n* = 33; Pearson *r* = −0.502, *p* = 0.002, *n* = 33). The dashed line indicates the linear regression fit with the shaded region indicating the 95% confidence interval. (B) Predicted CV increases significantly with impairment severity in simulations with increased muscle activity (Pearson *r* = 0.608, *p* = 0.016), but not in simulations with reduced muscle strength (Pearson *r* = 0.284, *p* = 0.347). The dashed lines indicate the linear regression fit with the shaded region indicating the 95% confidence interval.

## Discussion

### Impairment-driven changes in cost landscape curvature explain changes in stride speed variability after stroke

It has previously been proposed that cost and reward landscapes influence the amount of motor variability between trials [25, 74]. These ideas have been extended to human locomotion, where it has been demonstrated that humans dynamically adapt their movement strategy to constraints to optimize energetic cost during walking [27, 28]. While prior work has focused on healthy adults adapting to external perturbations, this framework is extended to the pathological case. Here, the concept of a dynamically changing cost landscape is extended to describe how neuromuscular impairments following stroke influence the cost landscape during locomotion and the variability of movement behavior.

The consistency between the predicted and observed coefficient of variation suggests that the cost landscape curvature captures a meaningful component of impairment-driven increases in stride speed variability after stroke. The simulations predict that increased paretic muscle activity will have a more significant effect on the coefficient of variation than reduced paretic muscle strength. However, the predicted coefficient of variation (3-14%) underestimates the overall coefficient of variation observed in the clinical post-stroke population (3-33%). This suggests that other mechanisms, such as increased sensorimotor noise, diminished feedback control, and muscle fatigue, are also highly relevant in describing the overall composition of stride speed variability observed after stroke. Similarly, variability has been shown to be introduced during the motor learning process as a means of discovering optimal strategies, suggesting that some fraction of variability can be attributed to exploration and exploitation during the learning [22, 75–77]. However, through trajectory optimization, it is assumed that the system is performing optimally during steady-state locomotion, thus it is “exploiting” the optimal form of each strategy after “exploring” alternative strategies through optimization.

The structure of the cost landscape, and specifically the sensitivity in cost to changes in speed, is proposed to influence the observed variability in stride speed despite the deterministic trajectory optimization simulations not capturing the effects of sensorimotor noise or feedback. The local curvature of the cost landscape, 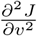, is analogous to mechanical stiffness, suggesting that cost landscape geometry regulates resistance to deviations from optimal speed in a way similar to mechanical impedance [78–80]. Under this interpretation, neuromuscular impairments that reduce the cost landscape curvature are analogous to a reduction in stiffness, thus permitting greater displacement from the optimal strategy. In the case of post-stroke locomotion, increased muscle activity signals in both the paretic and non-paretic limbs may lead to increased sensorimotor noise, which can contribute to observed variability [25, 31].

Under optimal feedback control frameworks, reduced cost sensitivity may reduce the drive for task-level error correction, since the penalty of suboptimal behavior becomes reduced compared to other effects [32]. Both increased sensorimotor noise and reduced need for task-level error correction provide explanations for the increase in variability observed after stroke not captured by a deterministic trajectory optimization model. This interpretation assumes that the feedback control system operates optimally in a restructured cost landscape, despite strong evidence that neural damage post-stroke could fundamentally alter the control system [72, 81–83].

### Stride speed variability as a biomarker of impairment-specific motor dysfunction

The relationship between cost landscape curvature and stride speed variability observed in this work suggests the utility of gait variability metrics as non-invasive computational biomarkers linked to changes in sensorimotor control and neuromuscular impairments that manifest after stroke. The simulations presented demonstrate that increases in paretic muscle activity are more strongly associated with increases in stride speed variability after stroke than reductions in muscle strength, but neither fully predict the magnitude of CV in a clinical post-stroke population. While stride speed coefficient of variation is strongly correlated with functional mobility scores, such as the Performance Oriented Mobility Assessment, changes in stride speed variability are influenced by a wide range of factors beyond the neuromuscular impairments investigated in these simulations. The results presented do suggest that stride speed coefficient of variation is sensitive to changes in motor control and strategy, without sensitivity to changes in muscle strength, suggesting the potential utility of CV as an indicator that can distinguish different forms of impairment. Overall, the correlation between stride speed CV and performance scores indicates the utility of using stride speed variability and other similar variability-based metrics as indicators of locomotor performance as a possible low cost and easily implementable metric that can be used to monitor changes in motor function.

### Limitations

Using a 2D musculoskeletal model for predictive simulations inhibits phenomena of stroke-like locomotion that appear in the medio-lateral direction, such as hip circumduction. The 2D model was chosen for this study because of its computational simplicity, which allowed for reliable solving in the hundreds of different optimization problems solved in the high-throughput framework. The 2D model is shown to capture the energetics of individual muscles reasonably well in the principal direction of movement, despite being constrained in the medio-lateral direction. Assessing the sensitivity of the system to the dimensions of the model and the initial conditions could help enhance the robustness of predictive simulations and expand their utility as a tool for understanding the underlying neuromusculoskeletal mechanics and the constraints that influence behavior. Using trajectory optimization to estimate the cost-minimized states for locomotion under simulated impairment has the additional limitation that it does not capture the effects of sensorimotor noise, which can have a large effect on observed variability and can be altered after impairment. Furthermore, stride speed variability was not simulated directly within the predictive modeling framework.

Trajectory optimization recovers deterministic cost-minimizing solutions and does not inherently model stochastic stride-to-stride fluctuations, which would require explicit noise formulations that scale non-linearly with simulation time. The relationship between cost landscape curvature and observed variability is therefore indirect, relying on theoretical assumptions about how the sensorimotor system responds to energetic gradients rather than on direct simulation of stride-to-stride fluctuations.

## Conclusion

This study demonstrates that neuromuscular impairments reshape the energetic cost of transport landscape for locomotion in ways specific to the type of impairment that influence the cost landscape curvature and stride speed variability. Increased paretic muscle activity flattened the local curvature and drove asymmetric non-paretic compensation more strongly than reduced paretic strength, even though both impairments lowered the cost-minimized walking speed and increased the cost of transport. Applying a relationship between cost landscape curvature and stride speed variability, it is predicted that the coefficient of variation increases significantly with muscle activity severity but not with reduced paretic muscle strength. Observed stride speed coefficient of variation is shown to exceed the model predictions suggesting that other mechanisms also influence overall stride speed variability. These findings provide a mechanistic account of impairment-driven increases in gait variability and suggest that stride speed variability carries information about impairment type that could support more targeted rehabilitation monitoring and assessment.

## Supporting information

**S1 Fig.**
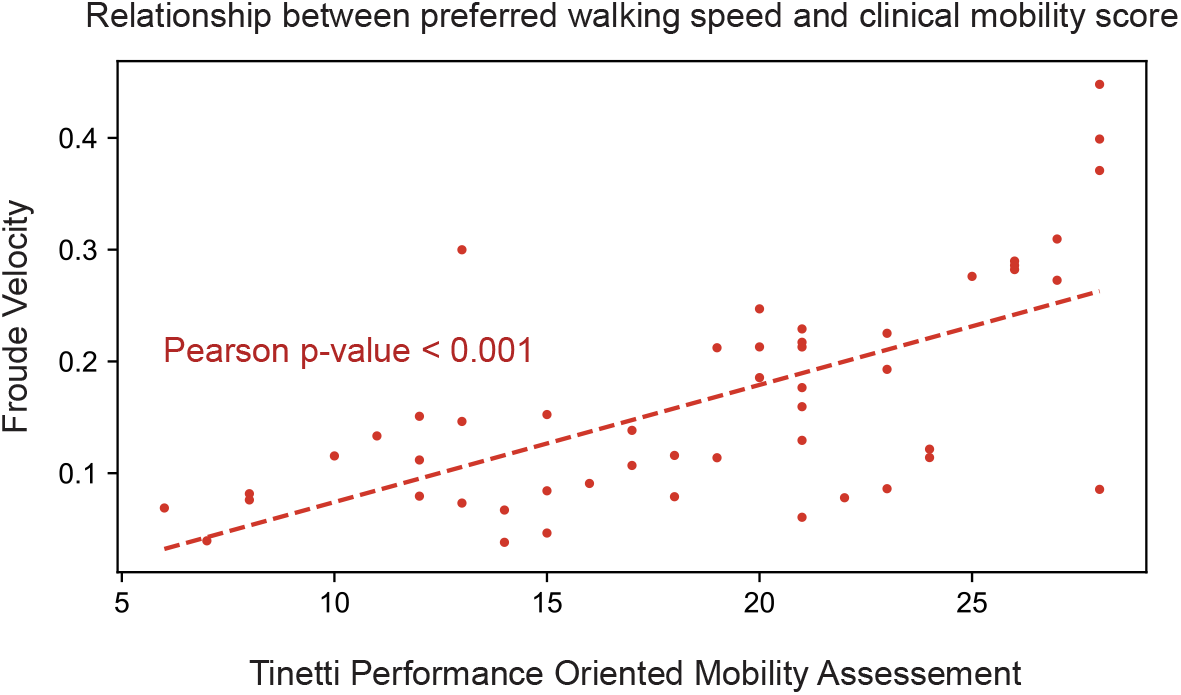
A statistically significant correlation is observed between the clinically assessed Tinetti Performance Oriented Mobility Assessment and the preferred walking speed in stroke survivors.

**S2 Fig.**
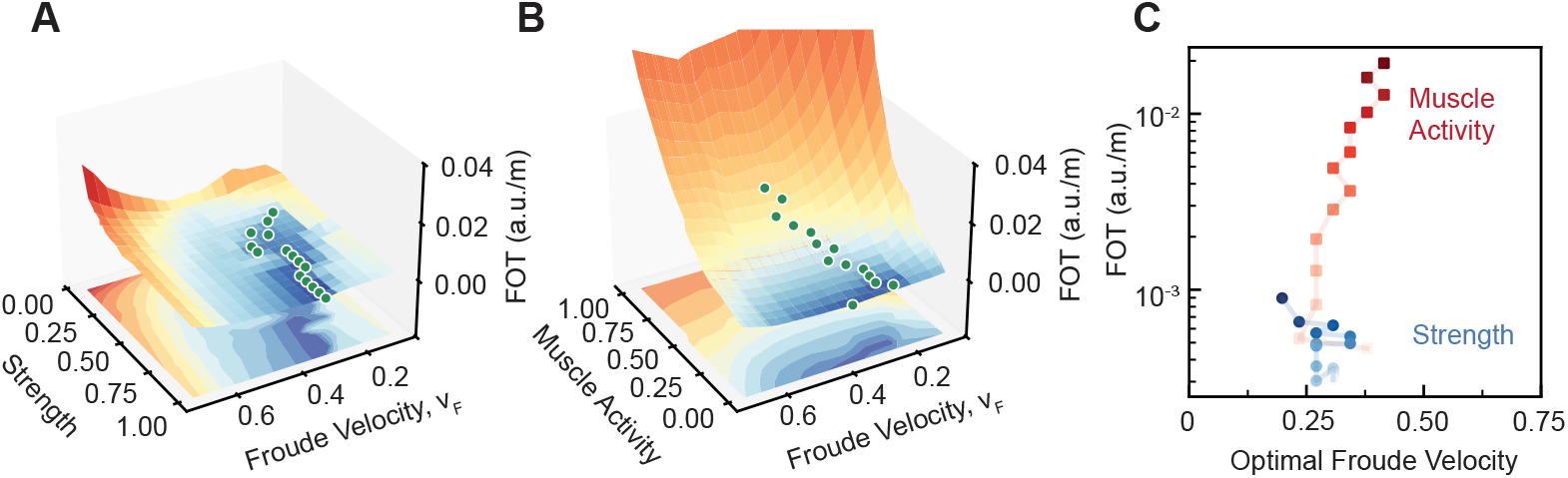
The fatigue cost of transport (FOT) reveals that both impairments reduce the cost-minimized speed. (A) The cost landscape of simulations with reduced muscle strength and the cost-minimized speed (green) show reductions in optimal speed. (B) The cost landscape of simulations with increased muscle activity and the cost-minimized speed (green) show reductions in optimal speed. (C) Comparing the cost-minimized speed between simulations with reduced muscle strength and increased muscle activity in the paretic limb reveals a maximum optimal speed of 0.4 and a minimum optimal speed of 0.2 for simulations with reduced muscle strength.

**S3 Fig.**
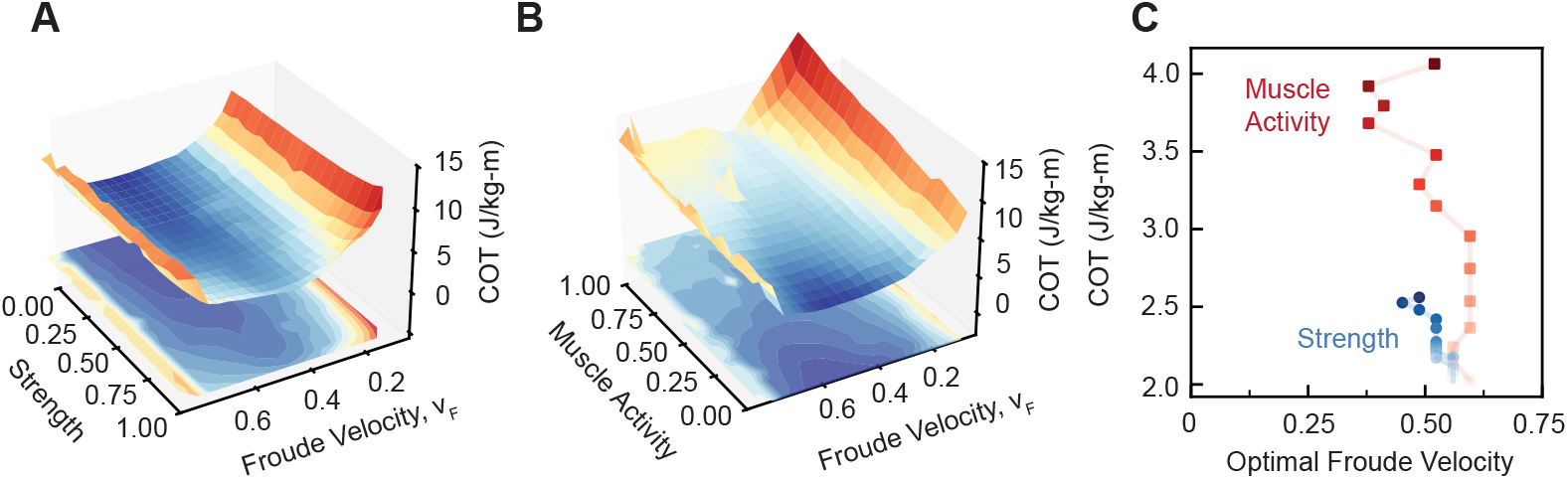
Metabolic cost-minimized speed was estimated in both sets of impairment. (A) Metabolic cost landscape for simulations with reduced paretic muscle strength. (B) Metabolic cost landscape for simulations with increased paretic muscle activity. (C) Metabolic cost-minimized speed for simulations with increased muscle activity (red) and reduced muscle strength (blue).

**S4 Fig.**
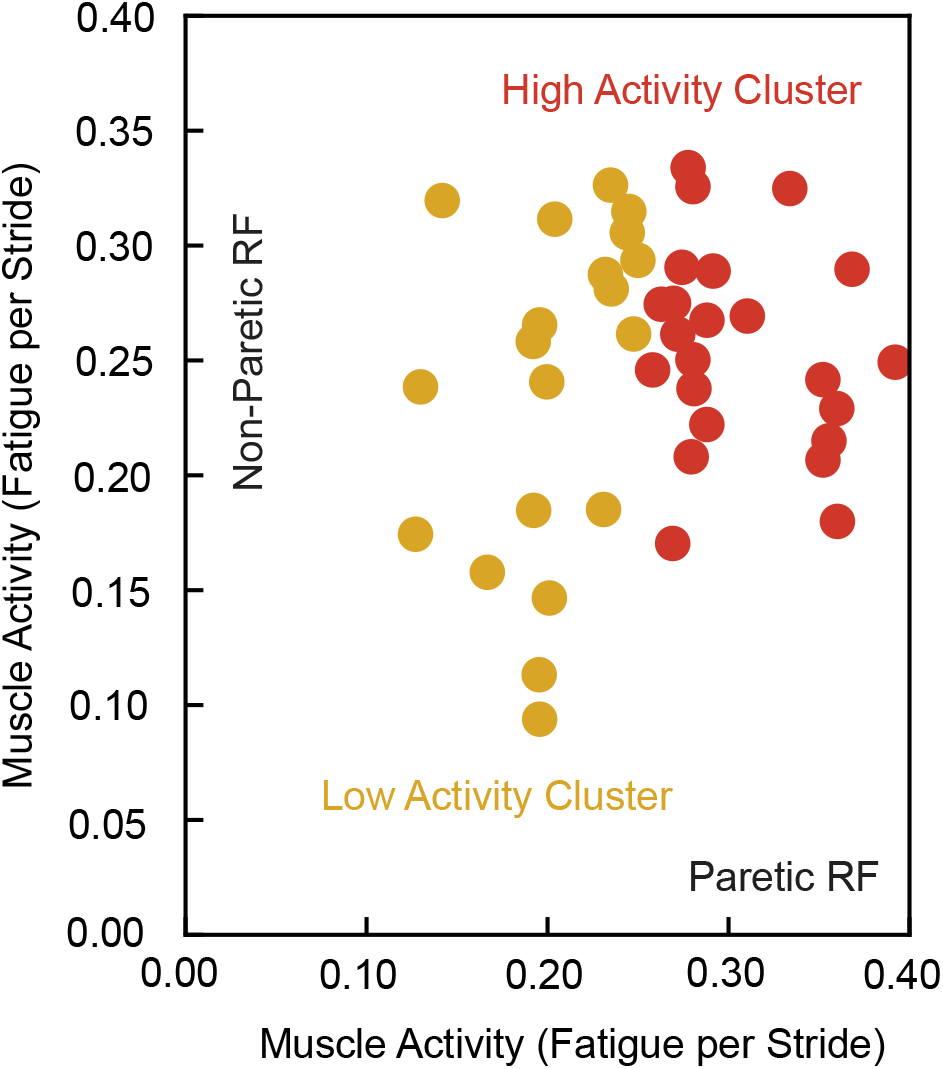
K-means clustering was applied in one-dimension to identify a group of stroke survivors with increased muscle activity in the paretic limb based on the magnitude of observed muscle activity in the paretic rectus femoris.

**S5 Fig.**
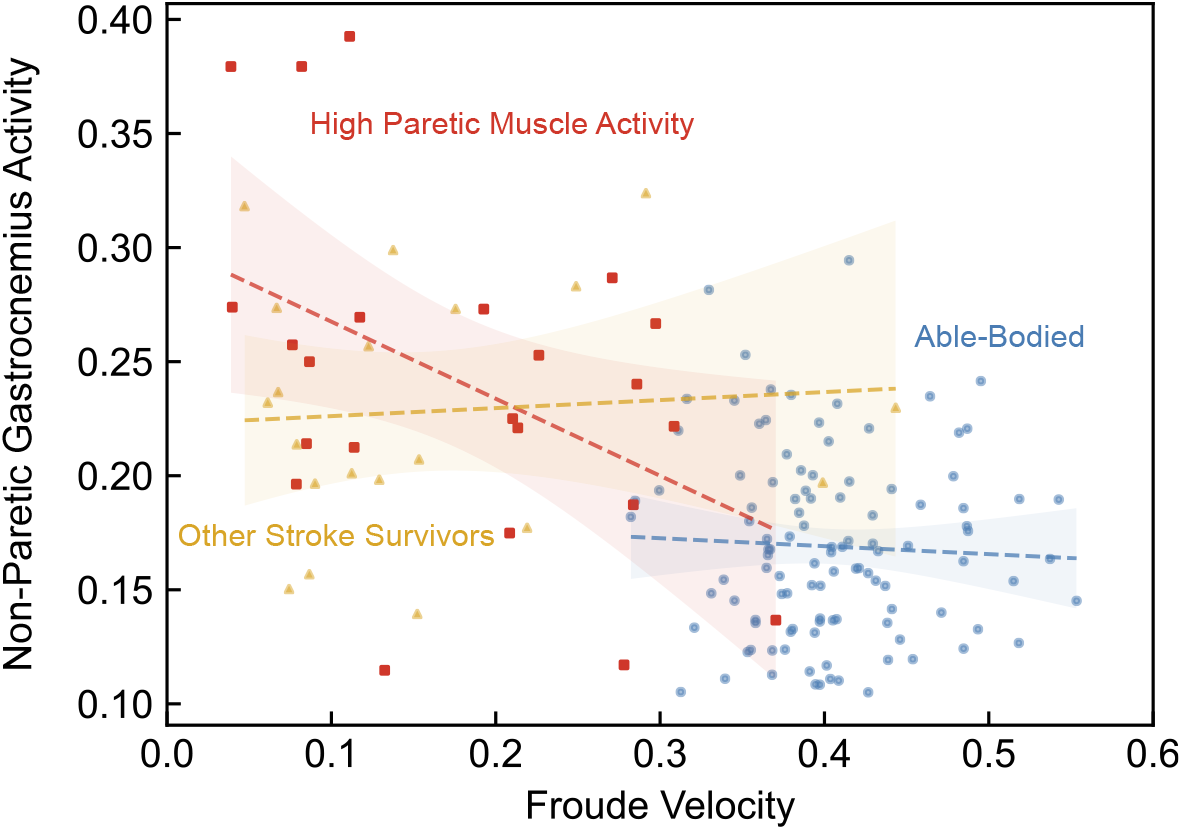
Subjects from the high muscle activity group are observed to have increased compensation from the non-paretic gastrocnemius as impairment severity increases. A statistically significant relationship between speed and non-paretic gastrocnemius activity is observed in the high-activity group (*p* = 0.031). In the low activity group and able-bodied group no significant correlations between speed and muscle activity in the non-paretic gastrocnemius are observed.

**S6 Fig.**
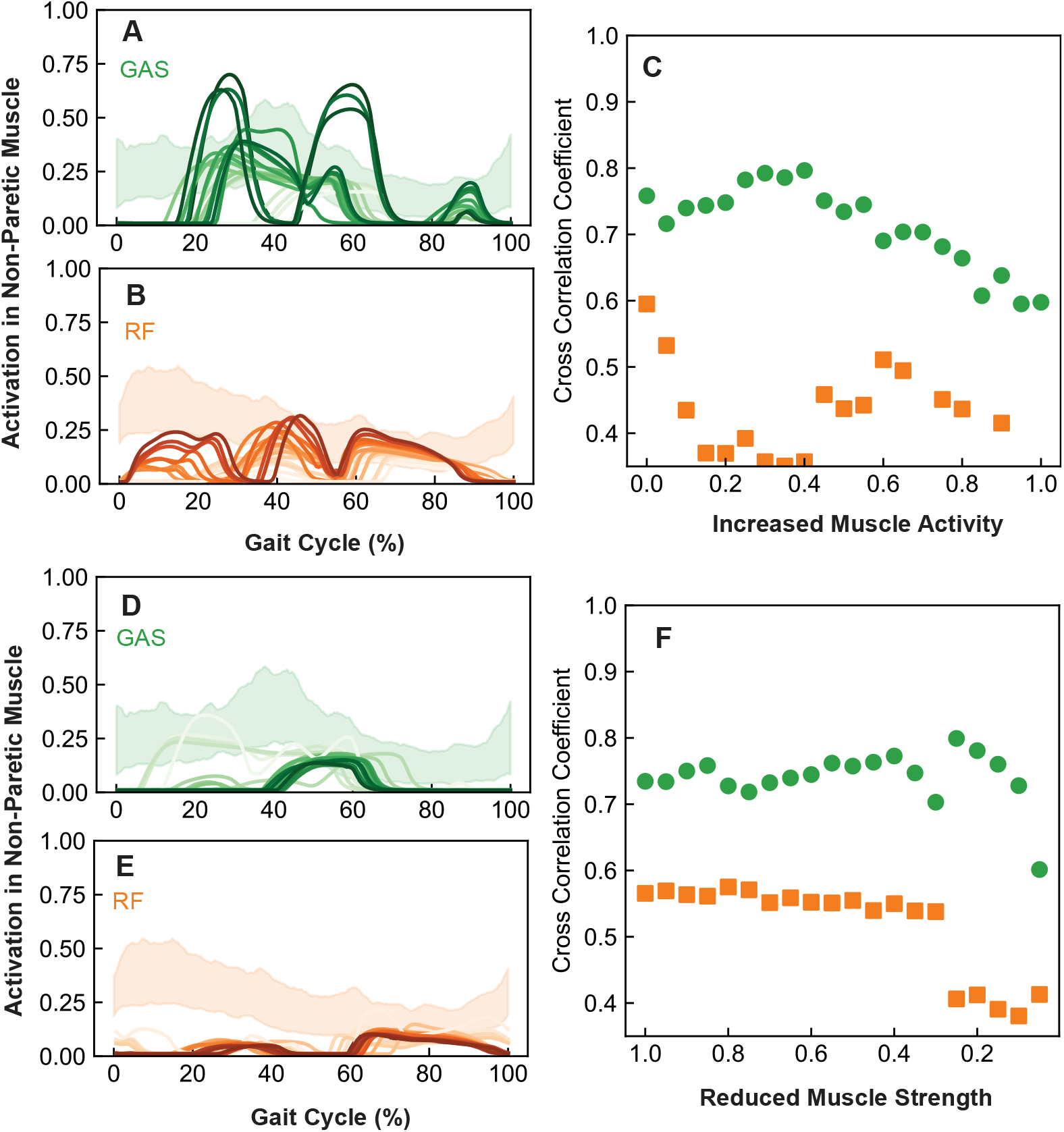
Simulated muscle activations through a gait cycle in the non-paretic limb are compared against the mean muscle activity profiles in stroke survivors. (A, B) The simulated muscle activations of the non-paretic gastrocnemius and rectus femoris at the optimal speed in simulations with altered muscle activity is compared against the standard deviation of the muscle activity profiles in stroke survivors at their preferred speed (shaded). (C) The max cross correlation coefficient is reported for each limb and increment of increased muscle activity in the paretic limb at the optimal speed. (D, E) The muscle activations for the gastrocnemius and rectus femoris at the optimal speed in simulations with reduced muscle strength are compared against the EMG muscle activity profiles in stroke survivors (shaded). (F) The max cross correlation coefficient is reported for each limb and muscle strength at the optimal speed.

**S7 Fig.**
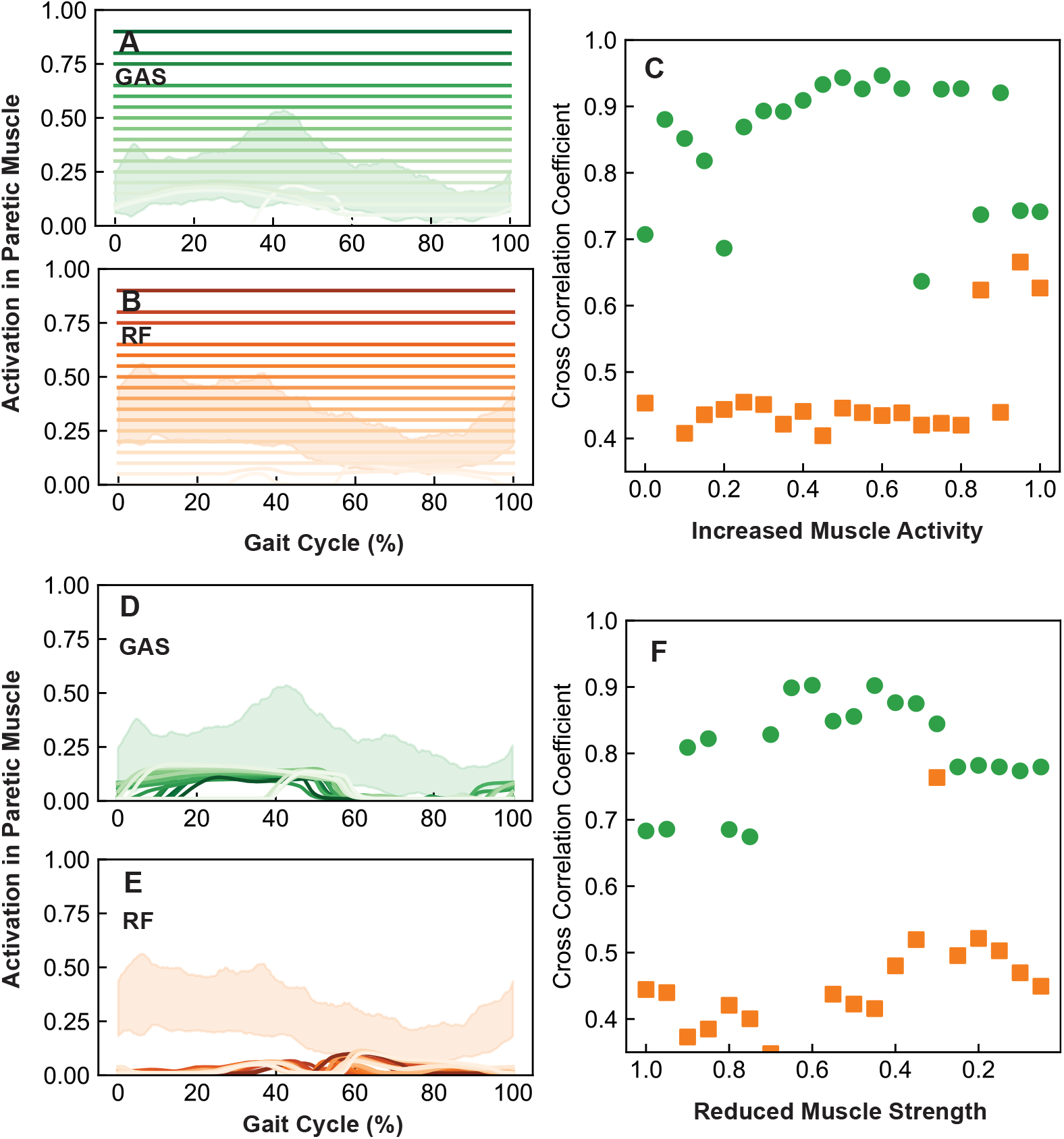
Simulated muscle activations through a gait cycle in the paretic limb are compared against the corresponding muscle activity profile in stroke survivors. (A, B) Simulated muscle activations of the paretic gastrocnemius and rectus femoris at the optimal speed in simulations with altered muscle activity are compared against the standard deviation of the muscle activity profiles in stroke survivors at their preferred speed (shaded). (C) The maximum cross correlation coefficient is reported for each limb and muscle activity at the optimal speed. (D, E) Muscle activations for the gastrocnemius and rectus femoris at the optimal speed in simulations with reduced muscle strength are compared with the real muscle activity profiles in stroke survivors (shaded). (F) The maximum cross correlation coefficient is reported for each limb and muscle strength at the optimal speed.

**S1 Appendix. Simplified relationship between the coefficient of variation and the cost landscape curvature**.

We postulate that changes in landscape curvature will influence the stride-to-stride fluctuations because small perturbations from the optimum speed will be increasingly costly when the curvature is higher (Fig. 1D). A simplified relationship and stride variability can be formalized by assuming that the sensorimotor system introduces motor variability that manifests as stride-to-stride fluctuations in speed that follow a Boltzmann-like distribution that is sensitive to relative changes in cost, such that

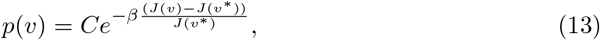

where *p*(*v*) is the probability the speed of a given stride and *C* and *β* are empirical constants. The Boltzmann-like distribution is motivated by theoretical and empirical work suggesting that the sensorimotor system is sensitive to relative changes in energetic cost during locomotion, such that movements incurring higher costs are less frequently expressed [28, 31, 60]. Assuming that near the optimal point the function can be expanded as a second-order Taylor series, the relationship between velocity and relative changes in cost becomes,

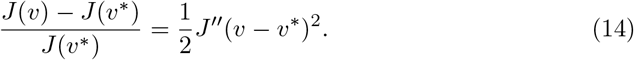

Then, it is recovered that stride-to-stride fluctuations would follow a Gaussian distribution,

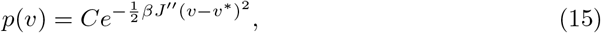

where the magnitude of the variance is linearly related to the curvature term,

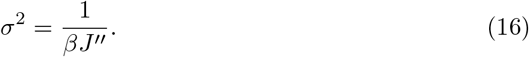

This results in the non-dimensional coefficient of variation being related to both the curvature and the optimal speed, such that

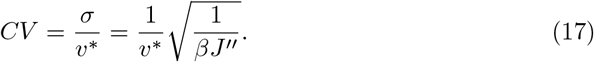

## Acknowledgments

This work was funded by the Sekisui House at MIT program in collaboration with the Institute for Medical Engineering and Science (IMES), the Center for Clinical and Translational Research at MIT, United States, and the Immersion Lab at MIT.nano. The authors acknowledge the MIT Office of Research Computing and Data for providing high performance computing resources that have contributed to the research results reported within this paper.

## Author Contributions

**Conceptualization:** Miles Smith, Praneeth Namburi, Nidhi Seethapathi

**Funding acquisition:** Brian W. Anthony

**Investigation:** Miles Smith

**Methodology:** Miles Smith, Nidhi Seethapathi

**Project administration:** Brian W. Anthony

**Resources:** Brian W. Anthony

**Software:** Miles Smith

**Supervision:** Brian W. Anthony

**Visualization:** Miles Smith

**Writing - original draft:** Miles Smith

**Writing - review & editing:** Miles Smith, Praneeth Namburi, Nidhi Seethapathi, Brian W. Anthony

## Notes

### Competing Interest Statement

The authors have declared no competing interest.

